# Functional ultrasound neuroimaging reveals mesoscopic organization of saccades in the lateral intraparietal area of posterior parietal cortex

**DOI:** 10.1101/2024.06.28.600796

**Authors:** Whitney S. Griggs, Sumner L. Norman, Mickael Tanter, Charles Liu, Vasileios Christopoulos, Mikhail G. Shapiro, Richard A. Andersen

**Author notes:** Correspondence should be addressed to W.S.G.

## Abstract

The lateral intraparietal cortex (LIP) located within the posterior parietal cortex (PPC) is an important area for the transformation of spatial information into accurate saccadic eye movements. Despite extensive research, we do not fully understand the functional anatomy of intended movement directions within LIP. This is in part due to technical challenges. Electrophysiology recordings can only record from small regions of the PPC, while fMRI and other whole-brain techniques lack sufficient spatiotemporal resolution. Here, we use functional ultrasound imaging (fUSI), an emerging technique with high sensitivity, large spatial coverage, and good spatial resolution, to determine how movement direction is encoded across PPC. We used fUSI to record local changes in cerebral blood volume in PPC as two monkeys performed memory-guided saccades to targets throughout their visual field. We then analyzed the distribution of preferred directional response fields within each coronal plane of PPC. Many subregions within LIP demonstrated strong directional tuning that was consistent across several months to years. These mesoscopic maps revealed a highly heterogenous organization within LIP with many small patches of neighboring cortex encoding different directions. LIP had a rough topography where anterior LIP represented more contralateral upward movements and posterior LIP represented more contralateral downward movements. These results address two fundamental gaps in our understanding of LIP’s functional organization: the neighborhood organization of patches and the broader organization across LIP. These findings were achieved by tracking the same LIP populations across many months to years and developing mesoscopic maps of direction specificity previously unattainable with fMRI or electrophysiology methods.

## Main

The posterior parietal cortex (PPC) integrates visual information with other sensory modalities, represents possible action plans, and decides upon the optimal action for downstream execution^1–3^. Separate PPC regions preferentially encode different movement types^4,5^, or “effectors”. Lateral intraparietal area (LIP) preferentially encodes saccades^6^, parietal reach region (PRR) preferentially encodes limb reaches^2^, and anterior intraparietal area (AIP) preferentially encodes grasping movements^7^. These areas reveal an effector-dependent functional organization within PPC.

It remains debated whether LIP possesses a mesoscopic functional organization for saccade directions^5,8^. Several studies have found that LIP’s response fields are spatially organized, with neighboring neurons having similar response fields^9–13^. However, the specific organization of these response fields remains debated^8^. This debate continues in part due to the limited field of view, sensitivity, and/or spatial resolution of existing recording techniques (**Fig. 1A**). fMRI can record whole brain activity, but lacks the spatial resolution and signal sensitivity to further refine our knowledge of LIP’s spatial organization (**Fig. 1B**). Intracortical electrophysiology can measure single neuron activity but cannot sufficiently sample or simultaneously record from large brain volumes, including primate PPC (**Fig. 1D**). Furthermore, it is difficult to align and reconstruct data recorded over many months, limiting our ability to observe anatomical patterns across many recordings. These limitations highlight the need for a sensitive technique to bridge the gap in spatial resolution between microscopic (e.g., single neurons) and macroscopic (e.g., whole brain) views of the primate cortex.

**Fig. 1.**
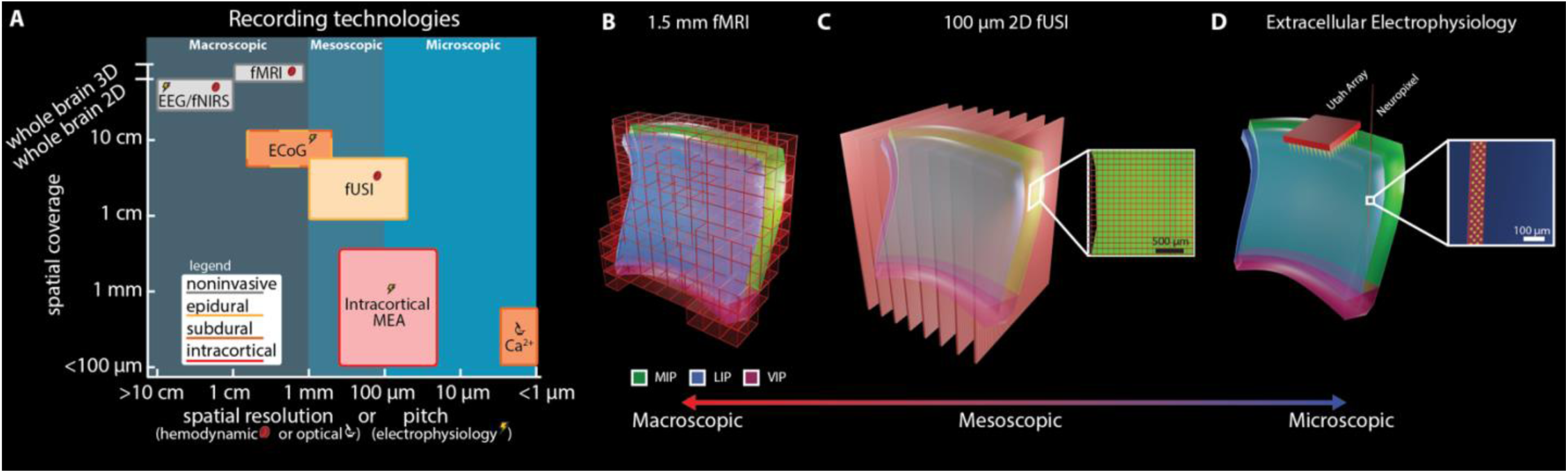
Functional ultrasound enables mesoscopic imaging of neural populations. **A.** Spatial coverage, invasiveness, and spatial resolution for different large animal recording technologies. Spatial coverage: Largest dimension of brain volume sampling. MEA: multi-electrode array; Ca^2+^: calcium imaging; ECoG: electrocorticography; EEG: electroencephalogram; fNIRS: functional near-infrared spectroscopy; fMRI: functional magnetic resonance imaging; fUSI: functional ultrasound imaging. Panel modified with permission from Griggs and Norman et al. 2024^17^. **B.** 1.5 mm isotropic fMRI. Each red box represents one voxel. **C.** 15.6 MHz 2D fUSI. Each red sheet represents one coronal imaging plane. Inset shows 100 μm x 100 x 400 μm voxel size. **D.** Utah array and Neuropixel 1.0 recording methods for recording from intraparietal sulcus. Inset shows size of Neuropixel 1.0 electrodes (yellow).

Here, we use an emerging technique, functional ultrasound imaging (fUSI) to determine the mesoscopic, i.e., between microscopic and macroscopic, spatial organization of saccadic response fields within LIP. fUSI’s large field of view, excellent sensitivity, and high spatial resolution (**Fig. 1A**, **C**) are ideally suited to this task^14–17^. We recorded fUSI while two rhesus macaque monkeys (Monkey L and P) performed an oculomotor task. We found functionally distinct subregions within (dorsal-ventral) and across (anterior-posterior) coronal LIP planes where small mesoscopic patches of neighboring cortex encoded different movement directions consistently across months to years. These results fill a gap in our understanding of LIP’s functional organization and demonstrate that fUSI is a powerful tool for elucidating mesoscopic function in the brain.

## Results

Using fUSI, we recorded high-resolution changes in cerebral blood volume (CBV) from multiple PPC subregions as two rhesus macaque monkeys (Monkey L and P) performed memory-guided saccades (**Fig. 2A**). These areas included lateral intraparietal area (LIP), ventral intraparietal area (VIP), medial intraparietal area (MIP), Area 5, Area 7, and medial parietal cortex (MP). During the task, each monkey fixated on a center cue, was cued with one of eight peripheral directions, remembered the cue location, and executed a saccade to the remembered location once the central fixation point extinguished.

**Fig. 2.**
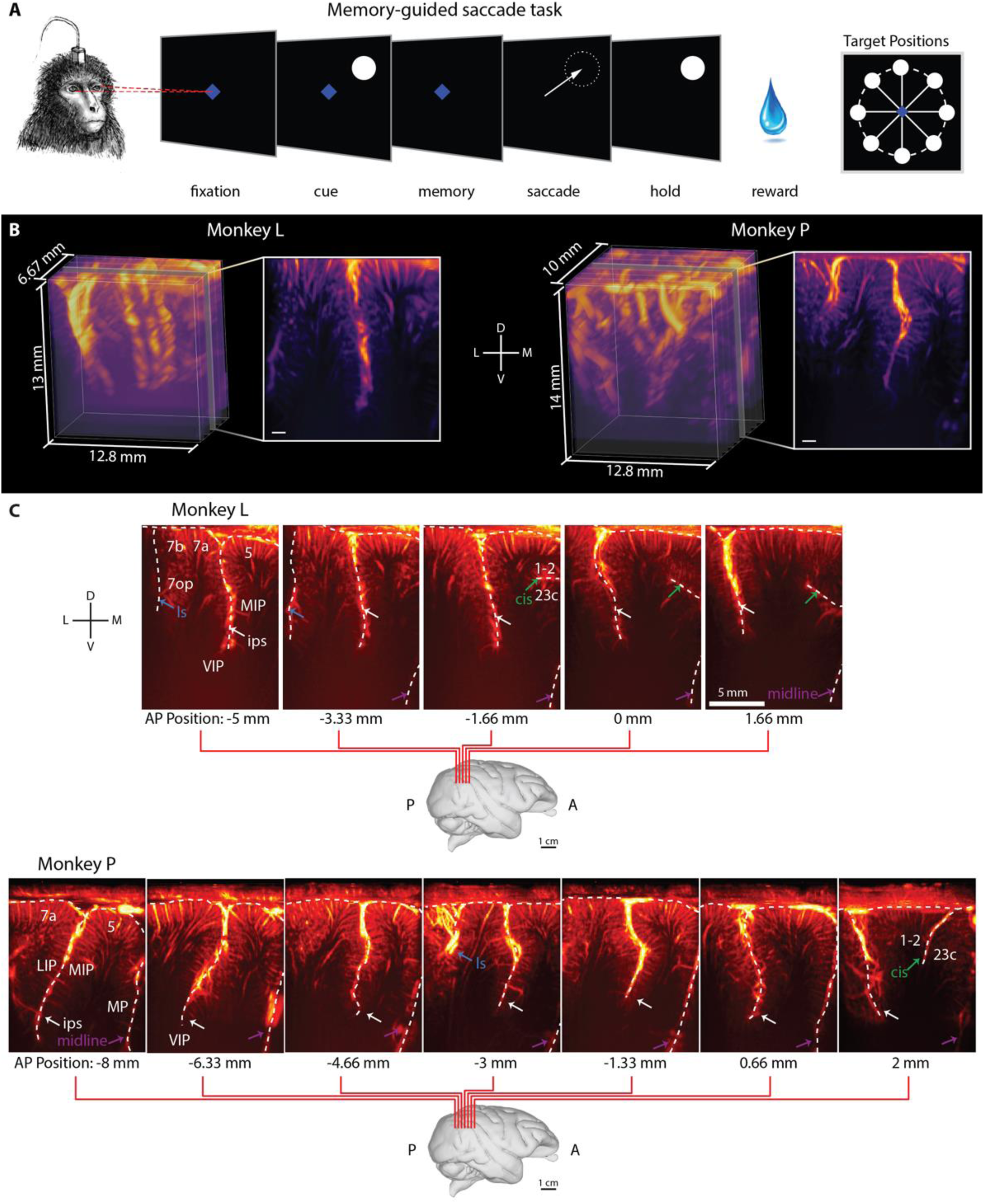
Monkeys performed memory-guided saccade task during fUSI acquisition. **A.** Memory-guided saccade task. A trial began with the monkey fixating on a center blue diamond. After the monkey fixated, a white circular cue was flashed in one of 8 peripheral locations. Once the center fixation diamond extinguished, the monkey made a saccade to the remembered cue location and maintained fixation on the peripheral location. If the saccade was to the correct location, the peripheral cue reappeared and the monkey received a liquid reward. **B.** 3D vascular maps for Monkey L and Monkey P. The field of view included the intraparietal sulci for both monkeys. White scalebar – 1 mm. D – dorsal. V – ventral. L – lateral. M – medial. **C.** Coronal imaging planes in Monkey L and P. Position relative to estimated ear-bar zero (EBZ) overlaid on a NHP brain atlas^18^. Anatomical labels based upon Saleem et al. 2012^19^. ls: lateral sulcus; cis: cingulate sulcus; ips: intraparietal sulcus.

We used a miniaturized linear ultrasound transducer array capable of high spatial resolution (100 μm x 100 μm in-plane) and a large field of view (12.8 mm width, 128 elements, 100 μm spatial pitch, 16 mm depth penetration, 400 μm plane thickness)^16,17^. We recorded 1 Hz fUS images by positioning the transducer surface normal to the brain above the dura mater. We recorded from multiple evenly-spaced coronal planes of the left PPC (**Fig. 2B, C**). We centered the recording chamber over the intraparietal sulcus to record from as much of the posterior parietal cortex as possible, both medial-lateral, but also anterior-posterior (**Fig. 2B**, **Supplemental Movie 1, 2**).

### Are there mesoscopic populations tuned to different directions?

To identify mesoscopic populations tuned to different directions, we used a general linear model (GLM) to identify voxels that responded differently to the eight directions. In both monkeys, mesoscopic PPC populations had clear directional tuning. In an example session from Monkey P (**Fig. 3A-C**), most LIP voxels (>75%) showed directional tuning while <4% of voxels outside of the LIP showed directionally-modulated activity. Some regions experienced substantial increases in CBV from baseline (>20%) where the magnitude of the increase depended on the direction. Other regions displayed suppression from baseline for certain movement directions (**Fig. 3B**). To quantify the example regions’ tuning, we averaged the response to each of the directions at the end of the memory period across all the voxels in each ROI to create regional tuning curves (**Fig. 3C**). Different populations within a single coronal plane had different preferred directions and different widths of their tuning curves. Some ROIs were broadly tuned to the entire contralateral hemifield (**Fig. 3B, C** – ROI 1) while other ROIs were tightly tuned to a narrow window of directions (**Fig. 3B**, **C** – ROI 2). The example session from Monkey L displayed similar phenomena (**Fig. 3D-F**). Most voxels (>70%) within LIP displayed directional modulation and these voxels were clumped into multiple subpopulations with different tuning curves. These regional tuning curves within LIP had different preferred directions and tuning curve widths. For example, ROI 1 displayed narrow tuning to 0° and 315° whereas ROI 2 displayed broader tuning to 45°, 90, and 135° (**Fig. 3D-F**). In Monkey L, the mid-MIP directly adjacent to the sulcus had some directionally tuned vascular response, but this activity did not penetrate deeper cortical layers. Including these MIP voxels, <4% of voxels outside of LIP showed directional modulation.

**Fig. 3.**
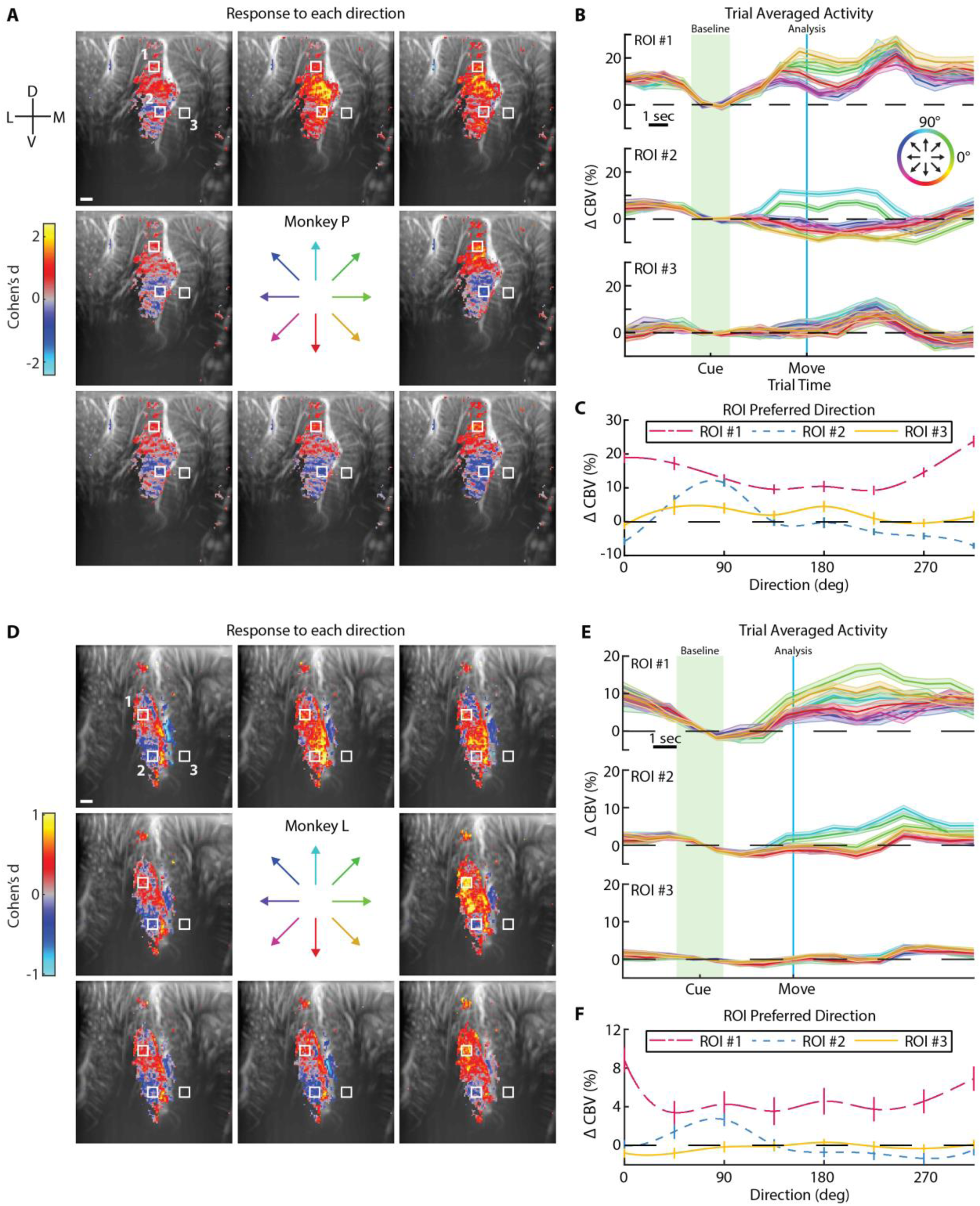
PPC contains multiple distinct directionally-tuned mesoscopic populations. **A.** Statistical parametric maps showing the average activity during the memory period. Voxel threshold determined by GLM F-test for voxels where q < 0.001 (FDR-corrected). White scale bar – 1 mm. Center arrows indicate the 8 directions tested. **B.** Event-related average of activity within each ROI. Each line represents one direction. The circular color scale indicates the direction of each line. Error shading shows SEM. Green shading shows timepoints used for calculating baseline and blue line shows timepoint used for analyzing memory response to the different directions. **C.** Tuning curves. Each line shows a cubic spline fit to the directional responses at the end of the memory period within each ROI. Error bars show SEM. **D-F**. Example session for Monkey L. Same format as **A-C**.

### How consistent is this directional tuning within a session?

Having observed clear mesoscopic populations with directional preference in both monkeys, we aimed to better understand the information content within these voxel subpopulations and the consistency of their responses from trial to trial. To this end, we performed decoding analyses for each example session. We trained a model to decode the intended movement direction on a subset of each example session’s trials and then tested how well the model could predict the intended movement direction on held-out test trials. If the model has statistically significant decoding accuracy on the test trials, it would demonstrate that the encoding of direction within the imaging field of view is consistent from trial to trial. For this decoding analysis, we used principal component analysis (PCA) to reduce the dimensionality of the fUSI data and linear discriminant analysis (LDA) to predict one of the eight movement directions using the PCA-transformed data. We examined the ability to decode the intended movement direction throughout the trial (**Fig. 4**) and found that we could begin decoding the intended movement direction significantly above chance (p<0.01; 1-sided binomial test) within 3 seconds of the directional cue onset (**Fig. 4A, D**). In both monkeys, the percent correct exceeded 50% (leave-one-out cross-validation; Monkey P – 59.6% correct, Monkey L – 54.1%). Missed predictions typically bordered the true movement direction (**Fig. 4B, E**). To quantify this, we present the mean absolute angular error between the predicted and true movement direction. As with the percent correct, the mean angular error reached significance (p<0.01; 1-sided permutation test) within 3 seconds of the directional cue. The mean angular error converged to <35° for both monkeys (Monkey P – 23.7°, Monkey L – 32.8°, **Fig. 4A-bottom, D-bottom**).

**Fig. 4.**
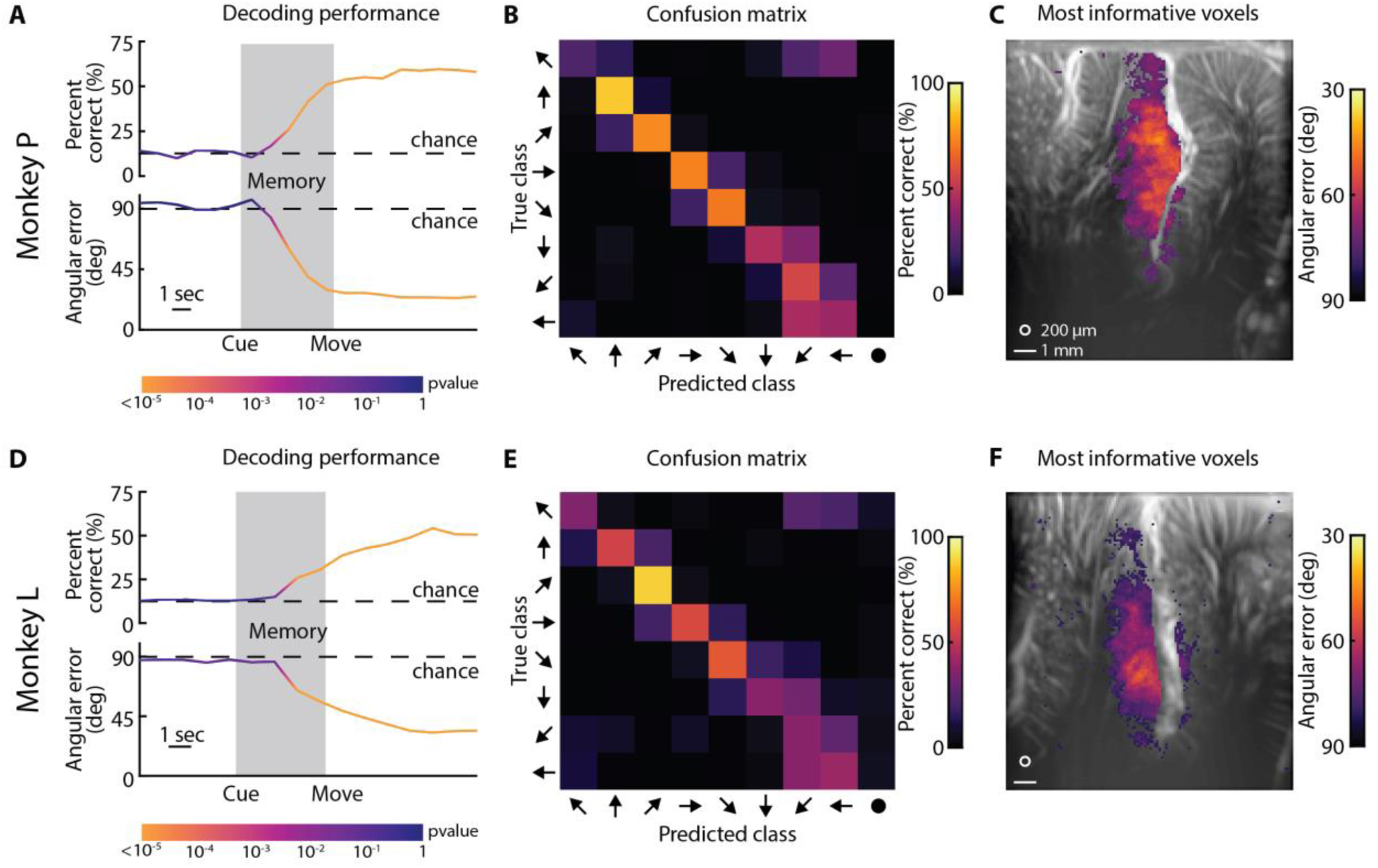
Single-trial decoding of eight intended movement directions with high accuracy. **A.** Decoding performance as a function of time. Top plot shows percent correct. Bottom plot shows mean angular error. Dashed lines show chance level performance. Color of the line shows statistical significance (1-sided binomial test or permutation test). **B.** Confusion matrix of decoding represented as percentage (rows adds to 100%). **C.** Searchlight analysis. Top 10% of voxels with the lowest mean angular error. White circle – 200 μm searchlight radius. White line – 1 mm scalebar. Masked voxels correspond to threshold of p<10^-5^. **D-F.** Decoding performance for Monkey L. Same format as A-C. Masked voxels corresponds to threshold of p<0.005.

We used the entire image at each timepoint to decode the intended movement direction on individual trials. One possibility is that the prediction is being driven by a few voxels that stay consistent while the other voxels fluctuate around. To understand which portions of the image, i.e., the vascular anatomy, contributed the most to the decoding performance, we performed a searchlight analysis. We moved a pillbox (200 μm radius) across the entire image and assessed the ability of each pillbox to decode the intended movement direction (**Fig. 4C, F**). In other words, we serially examine how each unique group of voxels contained within a 200-μm radius pillbox can individually decode the intended movement direction. To separate information contained within different brain regions, we only analyzed voxels on the same side of the sulcus for a given searchlight pillbox. As an example, a pillbox centered on an LIP voxel only contained LIP voxels whereas a pillbox centered on an MIP voxel only contained MIP voxels. We found that many of these 200-μm radius pillboxes, or voxel patches, could robustly decode the intended movement direction (p<0.01) with many of the voxel patches approaching 30° angular error (**Fig. 4C, F**). In other words, these searchlight results demonstrated that many PPC voxels encoded the intended movement direction and could drive accurate single-trial predictions. LIP contained the most informative voxels and these informative voxels overlapped with the same voxels identified with the previous GLM analyses (**Fig. 3A, D**). In both animals, a small number of voxels within MIP were within the 10% most significant voxels (threshold: Monkey L – p<0.005, Monkey P – p<10^-5^). In Monkey P, 7% of the most significant voxels were in MIP; in Monkey L, 9% of the most significant voxels were in MIP. These significant voxels within MIP were in the superficial cortical layers along the sulcus and did not extend into deeper cortical layers, matching the results from Monkey L’s GLM analysis.

### Are these mesoscopic populations stable across multiple days?

In the example sessions, PPC subpopulations were robustly tuned to individual movement directions, but is the function in each population stable across time? In a previous paper, we showed that populations in PPC could be used to control an ultrasonic brain-machine interface even after 60+ days since training the decoder model, suggesting that PPC populations are stable across at least 1-2 months^17^. To extend this result and better understand the stability across many months to years, we collected data from the same coronal plane across 4 – 30 months. We then trained our decoder on one session’s data and tested its performance on other sessions from the same plane without retraining or calibrating it. We tested all combinations of sessions. We hypothesized that, if the subpopulations’ functions were constant across time, a decoder trained on one session would accurately predict intended movement directions on another session’s data from the same coronal imaging plane.

In Monkey P, the decoder performed above chance level for over 100 days (**Fig. 5A**) and across all pairs of training and testing sessions (p<10^-5^; 36/36 pairs) (**Fig. 5B**). In Monkey L, the decoder performed above chance level for more than 900 days between the training and testing sessions (**Fig. 5D**), an effect that persisted across nearly all pairs of training and testing sessions (p<0.01; 117/121 pairs) (**Fig. 5E**). Cross-validated decoding performance varied within each training session (diagonal of performance matrices; **Fig. 5A, D**), so we also present cross-session decoding accuracy normalized to each training session’s cross-validated accuracy. We did not observe any clear differences between the absolute and normalized accuracy measures. Interestingly, in Monkey L, the decoder trained on the March 13, 2021 session performed the best for three directions (contralateral up, contralateral down, and ipsilateral down) in the training set and continued to decode these same three directions the best consistently throughout the test sessions (**Fig. 5D**). We saw this pattern where the decoder could best predict certain directions, even when the training session had poor cross-validated performance by itself (**Fig. S1**). We also observed in both monkeys that temporally adjacent sessions exhibited better performance (**Fig. 5C, F**).

**Fig. 5.**
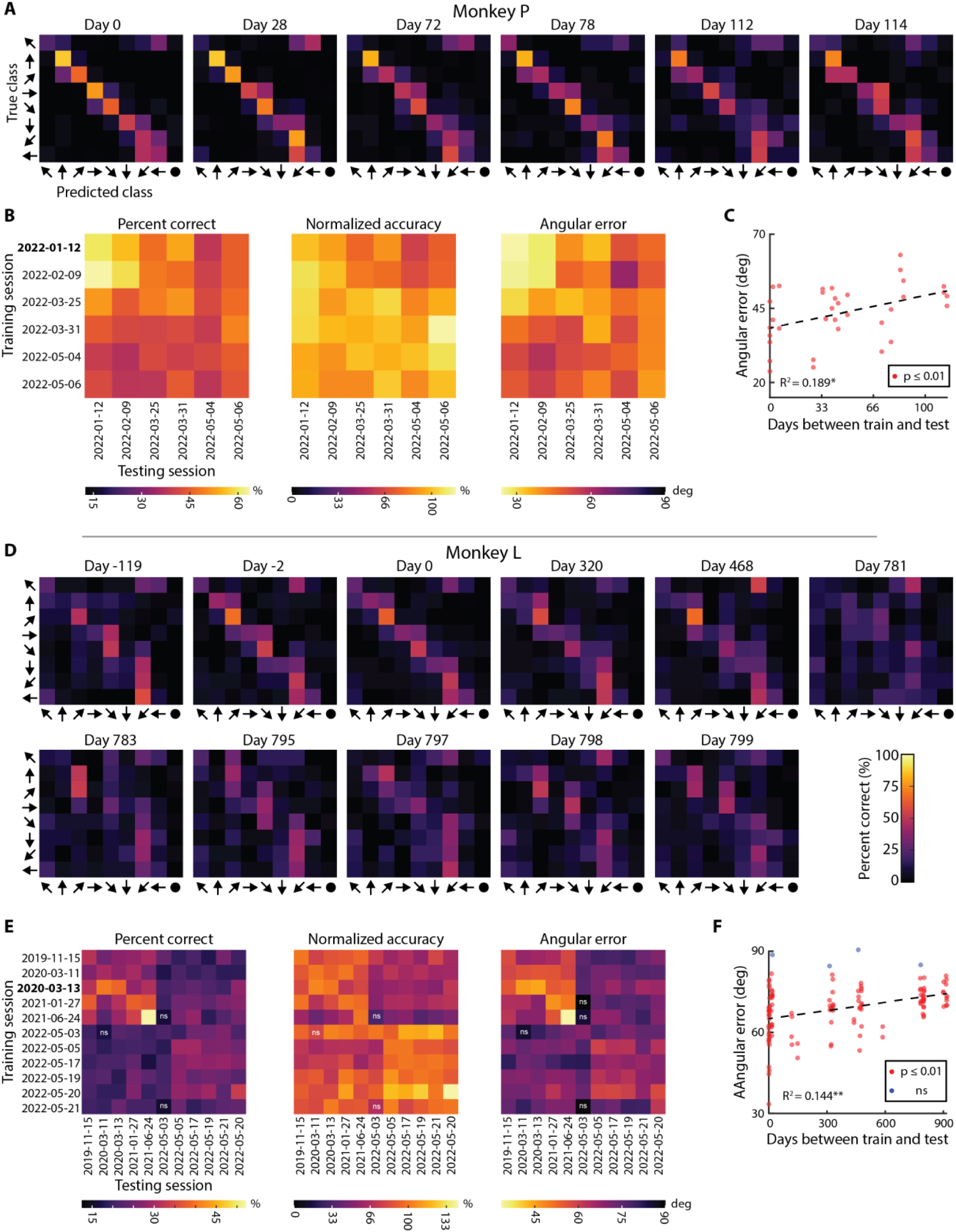
PPC stably encodes movement direction across many months to years. **A.** Example decoder stability for Monkey P. Trained the decoder on Day 0 data and tested the trained decoder on other sessions from the same imaging plane without any retraining. **B.** Decoder stability for training and testing on each session. ns – nonsignificant decoding performance (α = 0.01). Bold text represents example session shown in Fig. 5A. **C.** Mean angular error as a function of days between the training and testing session (absolute difference in time). Dashed line – Linear fit to data. *=p<10^-2^, **=p<10^-4^. **D-F**. Decoder stability for Monkey L. Same format as Fig. 5A**-C**.

In Monkey L, the performance was clumped into two temporal groups (before and after May 3, 2023). Was this change in performance due to physical changes in the imaging plane or due to changes in subpopulation function? Although we did our best to align our recordings to the exact same imaging plane from day to day, it is possible that alignment was imperfect and out-of-plane. Visual inspection revealed consistent macrovasculature, e.g. arteries, but inconsistent mesovasculature (**Fig. S2A, D**). Importantly, the physical changes would suggest that (a) we were decoding from slightly different neural populations and (b) small neighboring neural populations encode different directional information. To test our hypothesis that physical differences in imaging plane (and therefore differences in vascular anatomy) led to the decrease in decoder performance, we measured the similarity of the vascular anatomy across time using an image similarity metric: the complex-wavelet structural similarity index measure (CW-SSIM)^20^. The CW-SSIM clumped the vascular images into discrete groups (**Fig. S2B, S2E**), matching our qualitative assessment of image similarity. The similarity grouping also matched the pairwise decoding performance grouping in Monkey L (**Fig. 5E**). The decoder performance and image similarity were correlated (**Fig. S2C, F**). As image similarity decreased between the training session and each test session, so too did decoder performance. This supports our hypothesis that the decrease in decoder performance resulted from changes in the imaging plane rather than drift in each subpopulation’s tuning.

### How does mesoscopic population tuning change across anterior and posterior portions of PPC?

Having demonstrated that, within an imaging plane, there are PPC subpopulations robustly tuned to individual movement directions and these subpopulations’ tunings are consistent across many months to years, we next asked how direction tuning varied across different anterior to posterior coronal imaging planes. We repeated the same GLM analysis for data acquired from coronal planes evenly spaced throughout the PPC (**Fig. 2B, C, Supplemental Movies 1, 2)** and found the peak preferred direction for every voxel (**Fig. 6A**). Several patterns appeared. First, each coronal plane contained LIP voxels with directional modulation. Some of these planes contained large, contiguous patches of activity while other planes contained multiple discrete patches of activity. Second, each anatomical plane in both monkeys contained multiple LIP subpopulations with different tuning properties. These different subpopulations were sometimes in discrete patches and sometimes in the same contiguous patch. Third, posterior planes encoded more contralateral upward movements while anterior planes encoded more contralateral downward movements. Fourth, in Monkey P, regions outside of the LIP contained directionally modulated voxels, including in medial intraparietal (MIP), medial parietal (MP), and Area 5 cortex. We did not observe any activity within Area 5 of Monkey L and only observed very superficial activity within MIP of a single coronal plane (-3.33 mm of EBZ). Unfortunately, Monkey L’s chamber was more lateral and did not contain the same posterior portion of MP where we observed activity in Monkey P.

**Figure 6.**
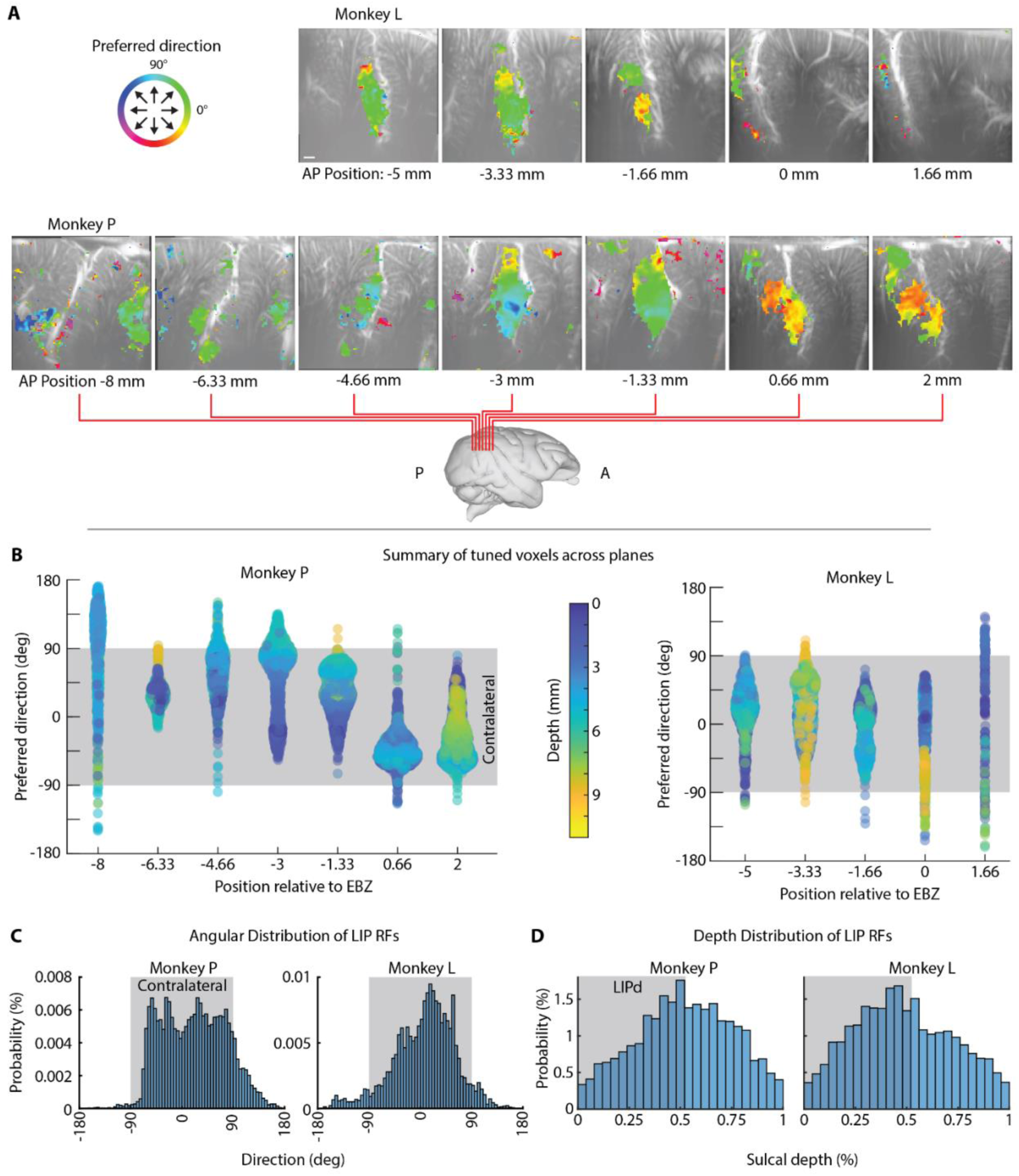
Polar direction is topographically organized along anterior-posterior axis of LIP. **A.** Color overlays showing preferred direction for voxels with statistically significant difference in response for different movement directions. Threshold based upon GLM F-test where p < 0.01 (FDR-corrected). **B.** Preferred direction for all significant voxels within each coronal plane. Color represents depth from brain surface. Grey shaded area shows contralateral angles. **C.** Angular distribution of response fields within LIP. Gray shaded area shows contralateral angles. **D.** Depth of tuned LIP voxels. Gray shaded area shows approximate LIPd.

To further understand the directional encoding across different coronal planes, we extracted the directional-modulated voxels and created a beeswarm chart containing each voxel’s directional preference (**Fig. 6B**). As expected, certain directions within a given plane were over-represented, i.e., clumps of similarly tuned neurons in the beeswarm plot. The anterior-posterior gradient was still evident where more anterior planes had more voxels tuned for downwards directions while posterior planes had more voxels tuned for upwards directions. Most voxels encoded for contralateral movements (-90° to +90°) although there were some voxels that responded most strongly for ipsilateral movements.

Each of the planes had broad and overlapping representation of contralateral movements. To better quantify these observations, we collapsed the voxels across planes (**Fig. 6C**) and found that >85% of tuned LIP voxels were contralateral preferring (Monkey P – 87.7%; Monkey L – 89.4%).

Certain anterior/posterior coronal planes had better representation of specific directions, so we asked whether there would be any performance difference in fUSI decoders trained on the different anatomical planes. This would have translational implications if specific anterior/posterior regions were better for directional decoding. We applied our decoding analysis to every recorded session (**Fig. S3A, B**). In Monkey P, all sessions reached statistical significance (18/18 sessions). In Monkey L, all but one session reached statistical significance (19/20 sessions). In Monkey P, the peak angular error within a session ranged from 17° to 55° (29.97° ± 2.32° mean ± SEM). In Monkey L, the angular error ranged from 33° to 85° (57.98° ± 3.35° mean ± SEM). There was no statistical difference (1-way ANOVA, α=0.01) between the percent correct or angular error depending on the plane (**Fig. S3C**). These results suggest that all anatomical LIP planes we sampled contained sufficient information to accurately decode at least eight intended movement directions on a single-trial basis.

Dorsal (LIPd) and ventral (LIPv) LIP are believed to have different functions with LIPd being involved in planning eye movements while LIPv is involved in both attentional and motor processes^21^. According to theories of topographic encoding^8^, we would expect separate representations of movement directions within LIPd and LIPv. We observed no clear split in function at the middle portion of LIP, so we relied upon a previous definition of 53% sulcal depth to compare activity within LIPd and LIPv^21^. We found that anterior LIPd was more active than posterior LIPd. Middle LIP, i.e., junction of defined LIPd and LIPv, consistently demonstrated the most activity across all planes. To quantify this observation, we labeled the beeswarm chart with the depth from brain surface of each voxel (**Fig. 6B**). We did not observe any clear trends in the data to clearly distinguish functional differences between LIPd and LIPv. We additionally collapsed all the tuned voxels across planes and looked at their percent depth within the sulcus (**Fig. 6D**). Instead of observing clear separation between LIPd and LIPv, we observed one homogenous group with the most activity peaking at the boundary between LIPd and LIPv, suggesting that LIPd and LIPv may share a topographic representation and/or that they receive common shared inputs.

## Discussion

Our results demonstrate that PPC contains subregions tuned to different directions. These tuned voxels were predominately within LIP and grouped into contiguous mesoscopic subpopulations. Multiple subpopulations existed within a given coronal plane, i.e., there were multiple preferred directions in each plane. A rough topography exists where anterior LIP had more voxels tuned to contralateral downwards saccades and posterior LIP had more voxels tuned to contralateral upwards saccades. These populations remained stable across more than 100 – 900 days.

### Sensitivity of fUSI

We observed large effect sizes with changes in CBV on the order of 10 –30% from baseline activity (**Fig. 3**). This is much larger than observed with BOLD fMRI where the effect size was approximately 0.4 – 2% on similar saccade-based event-related tasks^22,23^. Our results support a growing evidence base that establishes fUSI as a sensitive neuroimaging technique for detecting mesoscopic functional activity in a diversity of model organisms, including pigeons, rats, mice, nonhuman primates, ferrets, and infant and adult humans^15–17,24–31^.

### Anterior-posterior gradient

Several studies have reported a patchiness in direction selectivity with many neighboring neurons tuned to approximately the same direction followed by an abrupt transition to a patch of a different preferred direction^9,10,32^. These results match very closely with the results observed in this study where we found clusters within LIP tightly tuned to one direction with differently tuned clusters in close proximity within a given plane. The transition between patches with different directional tuning is spatially sharp and can occur in just a few voxels, with some transition zones lower than 500 µm. These results further emphasize the high spatial resolution of fUSI for functional mapping of neuronal activity. These results also closely match a previous study that used fUSI to identify the tonotopic mapping of the auditory cortex and inferior colliculus in awake ferrets where the authors found a functional resolution of 100 µm for voxel responsiveness and 300 µm for voxel frequency tuning^25^.

Our results support previous studies’ evidence of topography within LIP. This fUSI data (**Fig. 6**) matches two fMRI^11,12^ and two electrophysiology^10,33^ studies that found an anterior-posterior gradient where anterior LIP encodes for more downwards movements and posterior LIP encoded for more upwards movements Two studies^9,13^ found the opposite anterior-posterior gradient where anterior LIP encoded for upwards movements while posterior LIP encoded for downward movements. Two possible explanations for these contradictory findings have been proposed. Blatt et al. 1990^8^ suggested that there existed separate topographies for two of the main functions of LIP. One for attentional processing (anterior LIP – downward attention; posterior LIP – upward attention) and one for saccade planning (anterior LIP – upward saccades; posterior LIP – downward saccades). In the second possible explanation, Arcaro et al. 2011^12^ reconciled their fMRI data with the two electrophysiology studies by suggesting that the differences result solely because of differences in recording site location, i.e., the two electrophysiology papers recorded from different overlapping anterior-posterior ranges of LIP. Our combined range of recording agreed with the results that Arcaro et al. 2011 showed for visuotopic LIP (LIP_vt_) and caudal intraparietal cortex (CIP-2). Additionally, our results overlap with approximately 4 mm (-4 to -8 mm of EBZ) to the provided stereotactic zero, i.e., ear-bar zero (EBZ), in Blatt et al. 1990^9^. Over that range, we observed the same tuning of LIPv for contralateral upward saccades. Taken together, our results, using a memory-guided saccade task, disagree with the first interpretation and support the latter interpretation that differences in recording site location explain the difference in anterior-posterior gradients.

### Preference for contralateral space

Previous studies found that LIP responds strongest to contralateral stimuli and movements. At the single neuron level, approximately 80-90% of LIP neurons are tuned to contralateral directionsP^9,10^. Of note, Platt and Glimcher 1998^34^ reported no bias towards contralateral or ipsilateral in their recorded LIP neurons. At the macroscopic population level, the BOLD response in LIP is also almost exclusively contralateral preferring^11,12,22^. In the present study, we also found that LIP has strongly lateralized responses with ∼88% of LIP voxels preferred contralateral directions. The reasons for the apparent discrepancy with Platt and Glimcher 1998^34^ remain unclear.

### Differences between dorsal and ventral LIP

Previous studies found that peripheral targets were represented within the LIPv while foveal and parafoveal targets were represented within the LIPd^9–12^. In our study, we only tested a single eccentricity (20°) and observed activity within both LIPd and LIPv. In both monkeys, we observed less LIPd activity in the more posterior planes (< -3.33 mm of EBZ).

The overall distribution of tuned LIP voxels did not demonstrate a clear separation between LIPd and LIPv. Our study was not designed to interrogate the encoding of eccentricity, so future fUSI studies with foveal, parafoveal, and peripheral targets will be needed to explore the mesoscopic representation of eccentricity within LIP and PPC.

Previous studies have found that LIPd was primarily involved in oculomotor planning while LIPv contributed to both attentional and oculomotor processes^21^. Our task was aimed at understanding the oculomotor representation of different directions within LIP but was not designed to separate the effects of attention from oculomotor planning. In our study, we did not demonstrate a separation in directional representation between LIPd and LIPv with the most activity peaking at the boundary between LIPd and LIPv. This suggests that LIPd and LIPv may share a common topographic representation rather than having separate duplicated representations of angular direction.

### Directional saccadic activity outside of LIP

In Monkey P, we additionally observed directionally modulated activity outside of LIP in posterior-ventral MIP, Area 7, and medial parietal area (MP). MP has been previously identified as a saccade-related area in single-unit electrophysiology and fUSI studies^16,35^. Monkey L’s recording chamber was located lateral of MP areas. Nevertheless, our results in Monkey P, combined with observations in previous studies, support that MP may be an underexplored oculomotor planning region. The MP voxels preferred contralateral directions and did not display any clear organization of their response fields.

The posterior-medial MIP also contained directionally tuned activity. Some of this activity (-8 mm of EBZ) is in superficial cortical layers, perhaps reflecting inputs to MIP that relay directional information from upstream brain regions. We do not know why we see activity within the deeper layers of posterior-ventral MIP. Previous work found the functionally-defined parietal reach region (PRR), overlapping with the anatomical area MIP, responds predominately to reach movements^2,4,36^. Our task was a memory-guided saccade task with no reach component. Monkey P sat in an open chair with his hands and arms free while Monkey L sat in an enclosed chair with his hands and arms confined. Despite Monkey P being free to move his arms, we did not observe any arm movements related to the task itself. Future fUSI studies where we use a task with intermingled reaches and saccades will be useful in elucidating why we see saccade activity within MIP of one monkey.

More anterior portions of putative Area 7a displayed directionally tuned activity to contralateral movement directions. This is consistent with previous literature that found Area 7a neurons have visual receptive fields and display saccade-related activity^37–40^. We do not know why only a small region of Area 7A in Monkey P showed directional tuning or why we did not observe directionally tuned activity in Area 7a of Monkey L. One possibility is that the neurons within individual voxels of Area 7a display high heterogeneity in their response fields such that no consistent tuning appears at the mesoscopic population level.

### Stability across time

We demonstrated that we could decode intended movement direction using a decoder trained on data from a different session many months to years apart. This strongly suggests that the directional preference for the LIP subpopulations remains stable at the mesoscale. The decoder performed best when the training and testing sessions were close in time. We have three possible interpretations for this. First, the representations of direction within subpopulations drift in difficult-to-predict ways across time. Under this interpretation, we would expect that the decoders’ predicted movement directions would become increasingly random as more time elapses as the tuned voxels used for the model decorrelate. A second interpretation is that the subpopulations drift, but they drift at the same rate and in the same directions. This would lead to the tuned voxels staying correlated but encode for different directions. Under this interpretation, we would expect to see the decoder make increasingly more mistakes, but in a consistent manner. For example, the decoder might develop an error bias where instead of predicting the correct class, it consistently predicts a different direction in its place. A third interpretation is that the vascular placement relative to our recording plane changed across time (consistent with changes in the imaging plane) and that we are decoding from slightly different neural populations. Under this interpretation, the decoding errors should increase for neighboring directions because the tuned mesoscopic populations observed in this study extend in both anterior-posterior directions and smoothly transition to encoding different directions rather than having sharp transitions where neighboring voxels encode for completely different directions. This smooth transition means that the populations used for decoding will still be similar to the original populations being measured, consistent with changes in the imaging plane.

Our data best supports the third interpretation: the recording plane physically shifted over time. Rather than the errors becoming increasingly random as time progresses, the confusion matrices still had strong diagonal components, i.e., correct predictions, but with higher variance about that diagonal. Additionally, image similarity metrics drift across time, confirming that the imaging plane changed despite our best attempts (**Fig. S2**). Finally, the decoder performance and image similarity were positively correlated. This supports the interpretation that the subpopulations are stable across time with our decoder performance decreasing because of our imaging plane changing.

### Applications to ultrasonic brain-machine interfaces

We previously showed that we could decode movement timing (memory/not-memory), direction (contralateral/ipsilateral), and effector (hand/eye) simultaneously on a single-trial basis with high accuracy^16^. We recently also demonstrated that we could train monkeys to use a real-time fUSI brain-machine interface (BMI) for up to eight directions of eye movements^17^. Here, we extended these papers’ results in several aspects.

First, we demonstrated that we could achieve better decoding performance using offline recorded data (50-60% correct) than the accuracy reported for the online real-time fUSI-BMI data (∼38% correct). One explanation for this performance increase is motion correction. In the present study, we used *post hoc* motion-correction to minimize movement of the imaging plane across a session. In the real-time fUSI-BMI study, we did not implement motion correction. In the present study, we showed why mesoscopic populations are tolerant to a small amount of motion: similarly tuned voxels are more often spatially contiguous. However, even modest amounts of motion would alter the information available to the decoder, decreasing performance. One future method that may be well-suited to this problem is convolutional neural networks that can utilize local structure in images to maintain high performance rather than our existing decoder algorithms that assume features do not move across time.

Second, we demonstrated that we could decode above chance level with a static decoder model even after several years. In Griggs and Norman et al. 2024^17^, we collected data over 79 days, far fewer than the 900 days reported here. This suggests that future ultrasonic BMIs can constantly update an existing model rather than needing to be recalibrated daily. This is one current advantage of imaging-based BMIs over intracortical electrode-based BMIs. Intracortical electrode-based BMIs typically require frequent calibration or retraining due to their inability to record from the same neurons across multiple days^41^. By simply combining imaging-based BMIs with image alignment (2D plane and potentially 3D volume) to a previous session’s field of view, we can stabilize BMIs over long periods of time.

### Future studies

#### Record along intraparietal sulcus axis

In our study and most of the previous studies of LIP response fields, the topography changed along an anterior-posterior axis. Future studies could use 3D fUSI or align a 2D fUSI imaging plane along the intraparietal sulcus to acquire a larger anterior-posterior slice of LIP. This was not possible in the current animals due to the size of the ultrasound transducer and how our chamber was positioned off-axis relative to the intraparietal sulcus. This future study could significantly increase the longitudinal resolution compared to the current study and simultaneously improve effect sizes thanks to the ability to record anterior-posterior populations synchronously (in contrast to the current study that reconstructed these data over many sessions).

#### Eccentricity axis

Many studies have found a topography along an eccentricity axis with foveal and parafoveal targets being anterior of the peripheral targets representation^9–13^. In the current study, we presented our stimuli at a single eccentricity. Future studies could compare the representation of foveal, parafoveal, and peripheral targets within the LIP, potentially improving the field’s understanding of how angular direction and eccentricity are spatially organized. Exploring foveal representation of saccades may also require a different experimental task than used in this study due to the difficulties associated with tracking and measuring small saccades^42^.

#### Spatial autocorrelation of fUSI voxels

Each voxel (∼100 μm x ∼100 μm x ∼400 μm) contains approximately 65 neurons and 130 glia^43^, whereas each 1-1.5 mm^3^ fMRI voxel contains approximately 16,000-24,000 neurons and 32,000-48,000 glia. This suggests that fUSI can detect very local activity within neural circuits, including from within different cortical layers. However, fUSI measures changes in CBV and neurovascular coupling is complex^44–46^. Although every neuron within the brain is positioned within 15 μm of a blood vessel^47^, the contributions of different cell types are not sufficiently well understood to disentangle their contribution to the CBV signal. Additionally, neighboring voxels are supplied oxygen and nutrients by the same neurovasculature. This could contribute to an unknown extent to spatial autocorrelation of fUSI encoding (**Fig. S4**), confounding our ability to precisely identify the size and spatial separation of tuned populations. Motion of our imaging plane and spatial smoothing further increases this spatial autocorrelation.

There have been a variety of methods proposed for fMRI to handle the statistical consequences of spatial autocorrelation and calculate accurate statistical thresholds for cluster-wise inference^48–51^. However, to the best of our knowledge, no methods have been devised to separate the various contributors to the spatial autocorrelation, including correlated neuronal activity. Future experiments are required to disambiguate the contribution of correlated neurons versus other contributors to the size of neurovascular patches with similar tuning. Each voxel most likely contains neurons with a mixture of response fields with a bias towards specific response fields. Simultaneously recording fUSI signals and single neurons will be crucial for understanding the response properties within individual voxels and patches of similarly tuned voxels.

#### Directional tuning of cortical layers

Ultra-high field fMRI has enabled sub-millimeter voxel resolution and allowed researchers to study cortical layer-specific activity^52–54^, especially with CBV-based fMRI^55,56^. Similar laminar analyses are possible with fUSI because it measures CBV and has higher spatiotemporal resolution and sensitivity than UHF fMRI. To date, only one fUSI study has begun to explore this possibility. Blaize et al. 2020^57^ inferred cortical layer based upon cortical depth from the sulcus and found layer-specific ocular dominance within deep visual cortex. In the present study, we observed broad activity within LIP that did not appear to respect any laminar boundaries within the cortex. In both monkeys, we detected some directionally specific activity within the shallower layers of MIP (**Fig. 3D, 4C, 4F, 6A**). This may reflect activity within superficial input layers. We could qualitatively estimate the boundary between white matter and grey matter based upon the amount of organized mesovasculature observed in our vascular maps. However, the thickness of cortex varied within and across imaging planes, which prevented reliable estimates of cell layer based upon cortical depth. Future studies will be needed to better understand how to define cortical layers with fUSI, including studies to identify layer-specific properties detectable by ultrasound.

## Conclusion

Here, we used fUSI to demonstrate that the posterior parietal cortex (PPC) contains mesoscopic populations of neurons tuned to different movement directions. This organization changed along an anterior-posterior gradient and remained stable across many months to years. These results unify previous findings that examined the topographic organization of LIP at the macroscopic (fMRI) and microscopic (electrophysiology) levels. In one monkey, we additionally found robust saccade-related activity within the medial parietal (MP) cortex, a parietal area that warrants further investigation. Using the methods established here for tracking the same populations across many months to years, it will be possible to apply brain-machine interfaces and other technologies that are advantaged by stable recordings across time.

## Methods

### Experimental model and subject details

All training, recording, surgical, and animal care procedures were approved by the California Institute of Technology Institutional Animal Care and Use Committee and complied with the Public Health Service Policy on the Humane Care and Use of Laboratory Animals. We worked with two rhesus macaque monkeys (*Macaca mulatta*; 14-years old, male, 14-17 kg). Monkey L participated in two previous fUSI experiments^16,17^. Monkey P participated in one previous fUSI experiment^17^.

### General

#### Animal preparation and implant

We implanted a titanium headpost and custom square recording chamber on each monkey’s skull under general anesthesia and sterile surgical conditions. We printed or machined a 24 x 24 mm (inner dimension) chamber using Onyx filament (Markforged) for Monkey L and a similar chamber made of PEEK for Monkey P. We placed the recording chamber over a craniectomy centered above the left intraparietal sulcus.

#### Behavioral setup and task

Each monkey sat head-fixed in custom-designed primate chairs facing an LCD screen ∼30 cm away. We used a custom Python 2.7 software based upon PsychoPy^58^ to control the behavioral task and visual stimuli. We tracked their left eye position using an infrared eyetracker at 500 Hz (EyeLink 1000, Ottawa, Canada). Eye position was recorded simultaneously with stimulus information for offline analysis.

Monkeys performed a memory-guided saccade task (**Fig. 2A**) where they fixated on a center dot (fixation state), maintained fixation while a peripheral cue was flashed for 400 ms in one of eight locations (20° eccentricity, equally spaced around a circle), continued to maintain fixation on the center dot (memory state), and finally made a saccade to the remembered cue location (movement state). If they correctly made a saccade to the cued location, the peripheral cue was redisplayed and the monkey maintained fixation on the peripheral target until the liquid reward (30% juice; 0.35 mL monkey L and 0.75 mL monkey P) was delivered. To avoid the monkeys predicting state transitions, we used variable durations sampled from a uniform distribution for each task state. In Monkey L, the fixation and memory phase were 4 ± 0.25 seconds, the movement phase was 0.75 ± 0.15 seconds, and the intertrial interval (ITI) was 5 ± 1 seconds. For Monkey P, the fixation and memory phase were 5 ± 1 seconds, the movement phase was 1 ± 0.5 seconds, and the ITI was 8 ± 2 seconds.

### Functional ultrasound imaging

We used a programmable high-framerate ultrasound scanner (Vantage 256; Verasonics, Kirkland, WA) to drive the ultrasound transducer and collect pulse echo radiofrequency data. We used a custom plane-wave imaging sequence to acquire the 1 Hz Power Doppler images. We used a pulse repetition frequency of 7500 Hz with 5 evenly spaced tilted angles (-6° to 6°) with 3 accumulations to create one high-contrast compounded ultrasound image. We acquired the high-contrast compound images at 500 Hz and saved the images for offline construction of Power Doppler images. We constructed each Power Doppler image using 250 compound images acquired over 0.5 seconds. To separate the blood echoes from background tissue motion, we used an SVD clutter filter^59^. For more details on the functional ultrasound imaging sequence and Power Doppler image formation, please see Norman and Maresca et al. 2021^16^ and Macé et al. 2013^60^.

We used a 15.6 MHz ultrasound transducer (128-element miniaturized linear array probe, 100 μm pitch, Vermon, France). This transducer and imaging sequence provided us with a 12.8 mm (width) and 13-20 mm (height) field of view. The in-plane resolution was approximately 100 μm x 100 μm with a plane thickness of ∼400 μm. During each recording session, we placed the ultrasound transducer on the dura with sterile ultrasound gel. We held the transducer using a 3D-printed slotted chamber plug that minimized motion of the transducer relative to the brain. The slots were spaced 1.66 mm apart. This slotted chamber plug allowed us to acquire specific imaging planes across sessions. To help with later offline data concatenation, we acquired vascular maps using a single Power Doppler image and adjusted the transducer until the acquired vascular map closely matched a previously acquired vascular image for that chamber slot.

### Across session alignment and concatenation

We concatenated data across multiple sessions for each imaging plane. We first performed a semi-automated intensity-based rigid-body registration to align the vascular anatomy between sessions. As described above, during the acquisition, we minimized out-of-plane movement between sessions by matching each session’s imaging plane to a previously acquired template image for each chamber slot. See Griggs and Norman et al. 2024^17^ for more details.

### 3D visualization

We used MATLAB to export the vascular images to NIFTI format. We used Napari^61^, the ‘napari-niftì plugin^62^, and custom Python code to visualize the 3D reconstruction and save as images. The images were combined to form a movie using Da Vinci Resolve 17.4.4 Build 7 (Blackmagic Design).

### Quantification and statistical analysis

Unless reported otherwise, summary statistics are reported as mean ± SEM.

#### General linear model (GLM)

We applied several pre-processing steps before creating the GLM to explain the data. We first applied a Gaussian spatial filter (FWHM – 100 μm). We then applied a pixelwise high-pass temporal filter (1/128 Hz) to remove low-frequency drift. We finally used grand mean scaling to scale each voxel’s intensity to a common scale^63,64^. To build the general linear model, we convolved the regressors of interest with a hemodynamic response function (HRF). We used a single gamma function with a time constant (τ) of 1 second, a pure delay (δ) of 1 second, and a phase delay (*n*) of 3 seconds based upon a previous monkey event-related fMRI study^22^. The regressors of interest were fixation period, memory period, movement period, and reward delivery. For the memory and movement periods, we used separate regressors for each direction. We then fit the GLM model using the convolved regressors and scaled fUSI data. We used an F-test to identify voxels that had a statistically significant difference to the eight directions during the memory period.

#### Multiple comparison correction

For all voxel-wise p-values used and reported, we used false-discovery rate correction (FDR) to correct for the simultaneous multiple comparisons. This was implemented using MATLAB’s ‘mafdr’ function.

#### Preferred direction

We used a center-of-mass approach to find the preferred tuning of each voxel. For each voxel, we first calculated the Cohen’s d measure of effect size by comparing the response at the end of the memory period to the baseline (-1 to 1 seconds relative to cue onset). This gave us a standardized measure of response strength for each direction. We then scaled the peak response at each voxel to be 1. We then found the centroid for each voxel, which provided both a direction and magnitude. The direction represents the peak tuning direction while the magnitude represents the strength of that tuning. A value close to zero means no tuning while a value close to 1 means highly tuned to a specific direction. This method minimizes assumptions about shape of the response field, such as whether it is Gaussian. We then smoothed the resulting statistical map using a pillbox spatial filter (1-voxel radius).

### Within-session decoding analysis

Decoding intended movement direction on a single trial basis had five steps: 1) aligning the fUSI data and behavioral data, 2) preprocessing, 3) selecting data to analyze, 4) dimensionality reduction and class separation, and 5) cross-validation. First, we created the behavioral labels by temporally aligning the fUS data with the behavioral data. We could then label each fUSI timepoint with its corresponding task state and movement direction.

Second, we preprocessed the data by applying several operations. The first operation was motion correction. We used NoRMCorre to perform rigid registration between all the Power Doppler images in a session ^65^. We then applied temporal detrending (50 timepoints) and a pillbox spatial filter (2-voxel radius) to each Power Doppler image.

Third, we would then select what spatial and temporal portions of the data to use in the decoder model. We always used the entire image where each pixel is a single feature. We used a dynamic time window. At each timepoint before the cue, we used all timepoints since the start of the trial. For example, to test our ability to decode at 3 seconds after the trial start, we used the fUS images at 0, 1, 2, and 3 seconds. At each timepoint after the cue, we used all previous timepoints after the cue in the trial. For example, to test our ability to decode at 2 seconds after cue onset, we concatenated the data from 0, 1, and 2 seconds after the cue. We treated these timepoints as additional features in the decoder model. In other words, the input to our decoder model had N*T features, where N is the number of pixels in a single Power Doppler image and T is the number of timepoints.

Fourth, we split the data into train and test folds according to a leave-one-out or 10-fold cross-validation scheme. For the test sets, we stripped the behavioral labels. We then scaled the train and test splits by applying a z-score operation fit to the train data. We used the entire image for our features, i.e., each voxel’s activity was a single feature. To train the linear decoder on the training data, we used principal component analysis (PCA) for dimensionality reduction and linear discriminant analysis (LDA) for class separation. For the PCA, we kept 95% of the variance. For the LDA, we used MATLAB’s ‘fitcdiscr’ function with default parameters. We used a multicoder approach where the horizontal (left, center, or right) and vertical components (down, center, or up) were separately predicted and combined to form the final prediction. As a result of this separate decoding of horizontal and vertical movement components, “center” predictions are possible (horizontal — center and vertical — center) despite this not being one of the eight possible peripheral target locations. We then calculated the percent correct and absolute angular error for each sample in the test data.

Fifth, we then repeated the model training and testing for each consecutive fold of data. We finally found the mean accuracy metrics across all the folds, i.e., mean accuracy and mean angular error. To correct for testing the performance at every trial timepoint, we used a Bonferroni correction.

We used a 1-sided binomial test to calculate the p-values associated with the percent correct results and used a permutation test with 100,000 replicates to calculate the p-values associated with the angular error results. For the permutation test, each replicate was created by sequentially drawing X directional guesses from a uniform distribution of the eight possible directions, where X is the number of trials in the session. We then calculated the 1-sided p-value of each of our results by finding how many of the replicates were less than our observed mean angular error.

See Norman and Maresca et al. 2021^16^ and Griggs and Norman et al. 2024^17^ for more details on these methods.

### Across-session decoding analysis

To test whether we could use a decoder trained on a separate session’s data to decode movement intent in a different session, we applied the same steps as for the within-session decoding analysis with two differences. First, the training set was all the data from a specific session and the testing set was all the data from a different specific session. Second, to assess performance within the same train and test session, we used 10-fold cross-validation instead of leave-one-out cross-validation. The later sessions’ data (after March 25, 2022) are used in a previous publication and were acquired at a 2 Hz imaging rate with slightly different acquisition parameters. See Griggs and Norman et al. 2024^17^ for more details about the acquisition of these data. For the across-session decoding analysis, we down-sampled this 2 Hz data to 1 Hz to allow us to easily compare the same trial timepoints between the two sets of data.

### Image similarity

We compared the pairwise similarity of vascular images from different sessions by using the complex wavelet structural similarity index measure (CW-SSIM). The CW-SSIM quantifies the similarity of two images, where 0 is dissimilar and 1 is the same image^20^. We used the CW-SSIM over other forms of SSIM because it is more flexible in incorporating variations in image resolution, luminance change, contrast change, rotations, and translations. We used an implementation freely available from the MATLAB Central File Exchange^66^ with 4 levels and 16 orientations.

### Searchlight analysis

We defined a circular region of interest (ROI) and, using only the pixels within the ROI, we performed the within-session decoding analysis using 10-fold cross-validation. We assigned that ROI’s percent correct and angular error metrics to the center voxel. We then repeated this across the entire image, such that each image pixel is the center of one ROI. To visualize the results, we overlaid the performance metric (angular error or percent correct) onto a vascular map and kept up to the 10% most significant voxels. As part of this searchlight analysis, we ignored activity within the sulcal fold or activity on the other side of the sulcal fold. To do this, we defined the boundaries of the sulcal folds using a custom GUI in MATLAB and only used voxels on the same side of the sulcal fold as the searchlight center. This is similar in principle to the cortical surface-based searchlight decoding developed for fMRI^67^.

### Spatial autocorrelation

For every pixel in the image, we examined voxels at different distances from the seed voxel. For each distance tested, we identified voxels that were between [max(0, i-0.1) mm, i mm] away. We then performed Pearson linear correlation between these identified voxels and the seed voxel. We then assigned the mean correlation to the seed voxel. To calculate the mean correlation for each distance, we took the mean and standard deviation across the entire image.

## Common abbreviations used

CBV: Cerebral blood volume
fUSI: Functional ultrasound imaging
GLM: General linear model
ips: Intraparietal sulcus
LDA: Linear discriminant analysis
LFP: Local field potential
LIP: Lateral intraparietal area
MIP: Medial intraparietal area
MP: Medial parietal cortex
PCA: Principal component analysis
PPC: Posterior parietal cortex
PRR: Parietal reach region
ROI: Region of interest
VIP: Ventral intraparietal area

## Acknowledgments

We thank Kelsie Pejsa for assistance with animal care, surgeries, and training. We thank Tyson Aflalo, Claire Rabut, Ernesto Criado Hidalgo, and Geeling Chau for their helpful discussions and insights. W.S.G was supported by an NEI F30 (NEI F30 EY032799), the Josephine de Karman Fellowship, and the UCLA-Caltech MSTP (NIGMS T32 GM008042). S.L.N was supported by the Della Martin Foundation. This research was supported by the National Institute of Health BRAIN Initiative (grant 1R01NS123663-01 to R.A.A., M.G.S., and M.T.), the T&C Chen Brain-Machine Interface Center, and the Boswell Foundation (R.A.A.).

## Author contributions

W.S.G., S.L.N., V.C., M.G.S., and R.A.A. conceived the study; W.S.G. trained the NHPs and acquired the data; W.S.G. and S.L.N. performed the data processing and analysis; W.S.G. drafted the manuscript with substantial contributions from S.L.N, M.G.S, and R.A.A., and all authors edited and approved the final version of the manuscript. V.C., C.L., M.T., M.G.S., and R.A.A. supervised the research.

