## Supplemental Figures for "Functional ultrasound neuroimaging reveals mesoscopic organization of saccades in the lateral intraparietal area of posterior parietal cortex"

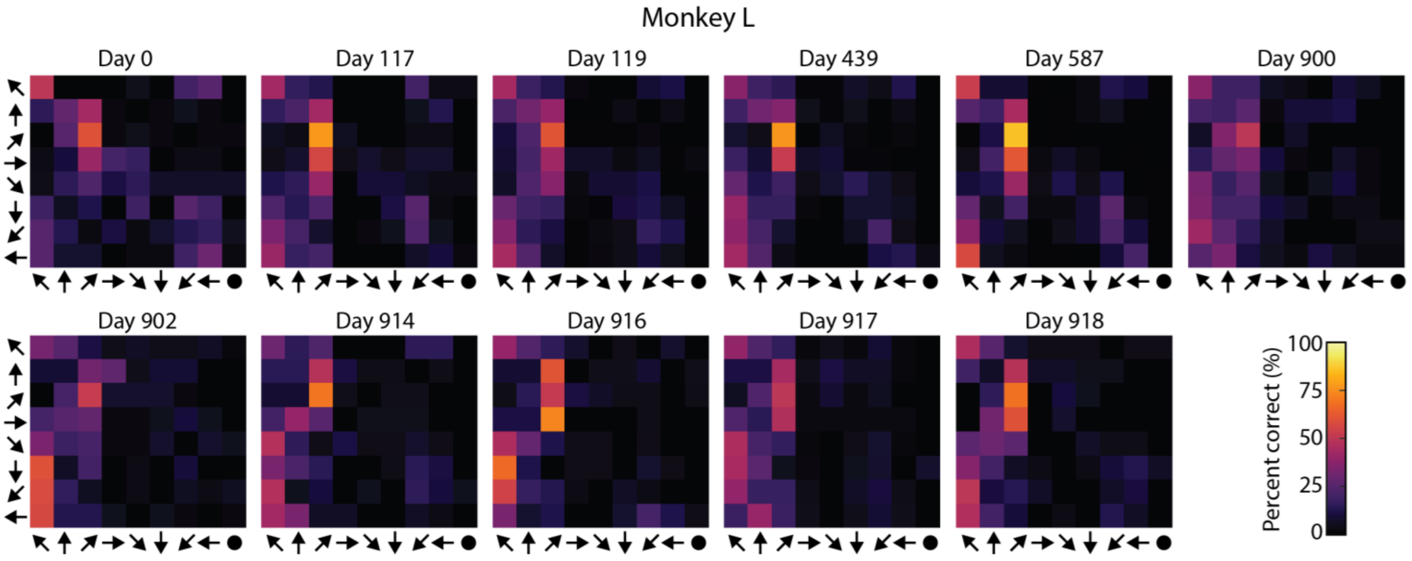


***Fig. S1 – Stability of directional preference across time.***
*Same format as **Fig. 5D** but using a different training session. The decoder was trained on Day 0 data and tested on other sessions from the same imaging plane without any retraining.*

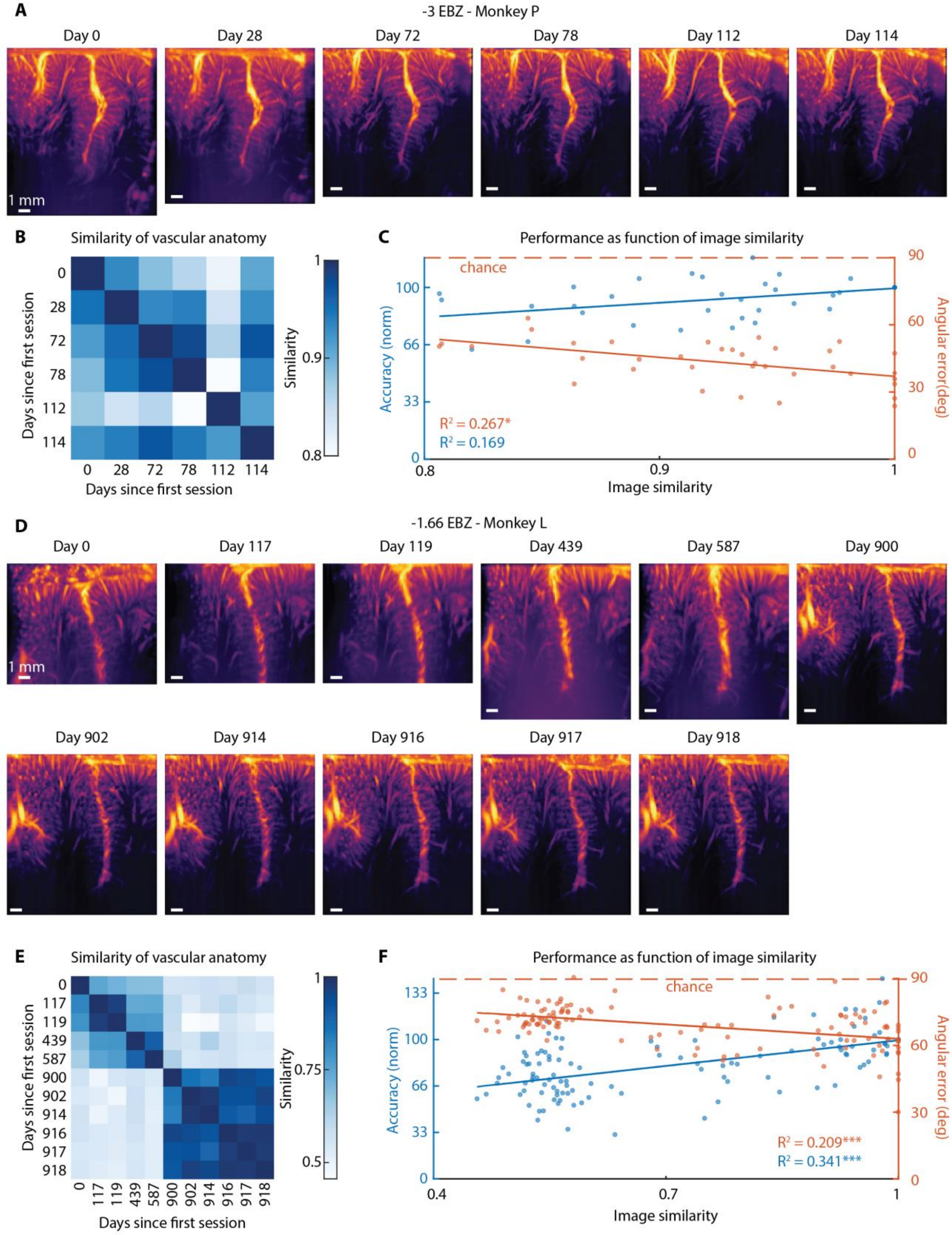


***Fig. S2 – Impact of image similarity on decoder performance.***

***A.*** *Vascular anatomy for each recording session from the same recording slot in Monkey P. White scalebar – 1 mm.*

***B.*** *Pair-wise similarity between different vascular images for Monkey P.*

***C.*** *Performance as a function of image similarity for Monkey P. Left axis (blue) shows normalized accuracy. Right axis (orange) shows mean angular error. Each session is represented by a pair of blue and orange dots.* *=$p<10^{-2}$, **=$p<10^{-4}$, ***=$p<10^{-6}$.

***D-F.*** *Same format as* ***(A-C)*** *for Monkey L.*

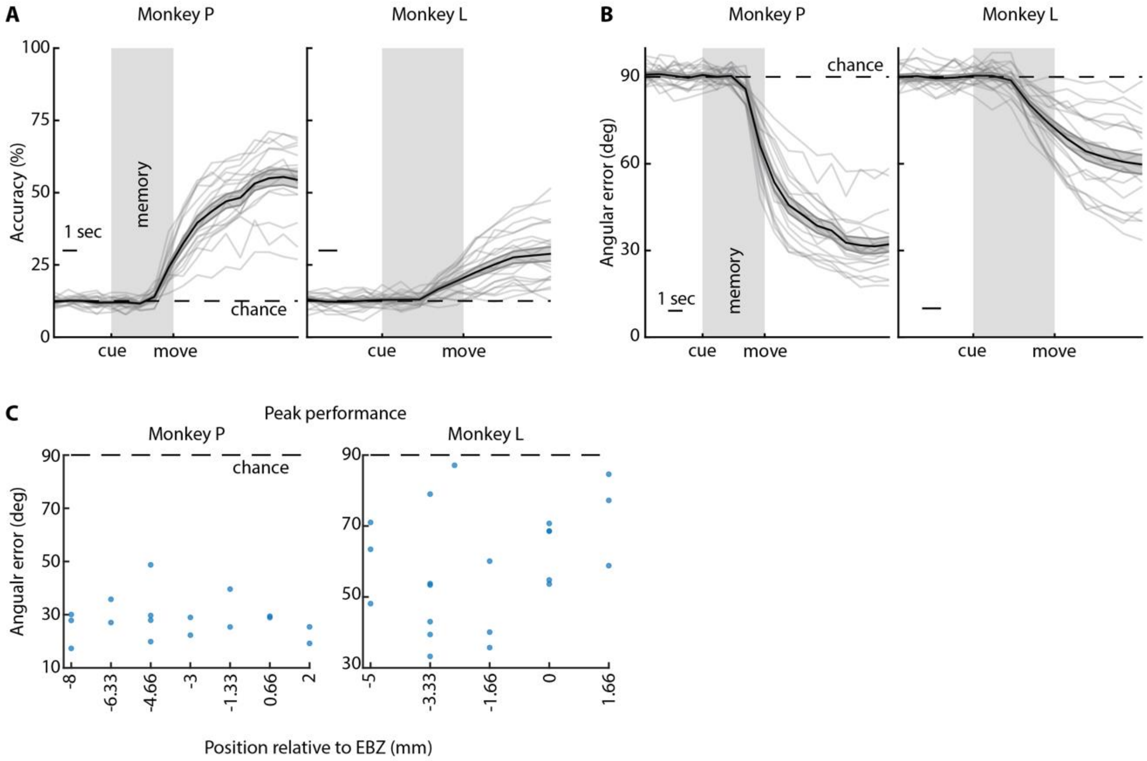


***Fig. S3 – Linear decoders can decode intended movement direction from most fUSI sessions, regardless of PPC plane.***

***A.*** *Percent correct for each session. Solid black line with gray envelope show mean ± SEM. Each gray line shows performance on single session. Dashed line shows chance level.*

***B.*** *Mean angular error for each session. Same format as in (A).*

***C.*** *Mean angular error as function of coronal plane.*

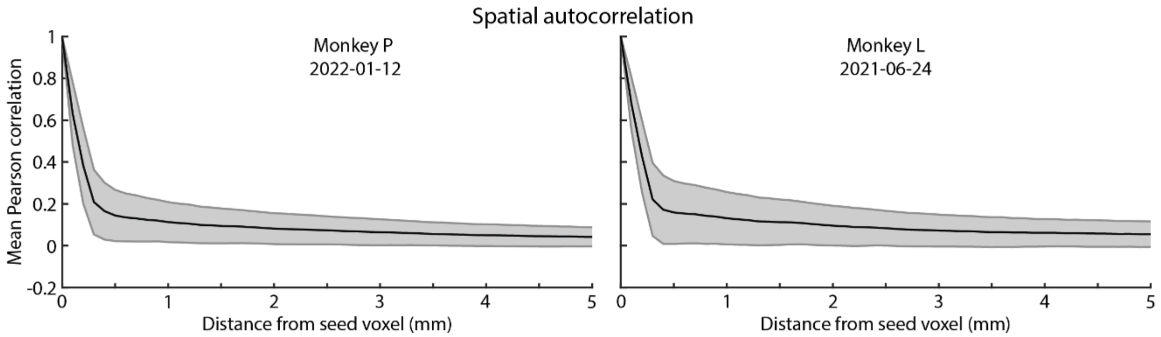


***Fig. S4 – Power Doppler data displays spatial autocorrelation.***

*Mean Pearson spatial autocorrelation across entire fUSI field. Voxel radius – Distance of voxels from center seed voxel. Shaded area – standard deviation*

### SUPPLEMENTAL MEDIA

**Supplemental Movie 1 Anatomy in Monkey P -** 3D reconstruction of vascular anatomy in Monkey P.
**Supplemental Movie 2 Anatomy in Monkey L -** 3D reconstruction of vascular anatomy in Monkey L.
